# The maize *PLASTID TERMINAL OXIDASE* (*PTOX*) gene controls the carotenoid content of kernels

**DOI:** 10.1101/2023.03.13.532334

**Authors:** Yongxin Nie, Hui Wang, Guan Zhang, Haiping Ding, Beibei Han, Lei Liu, Jian Shi, Jiyuan Du, Xiaohu Li, Xinzheng Li, Yajie Zhao, Xiaocong Zhang, Changlin Liu, Jianfeng Weng, Xinhai Li, Xiansheng Zhang, Xiangyu Zhao, Guangtang Pan, David Jackson, Qingyu Wu, Zhiming Zhang

## Abstract

- Carotenoids perform a broad range of important functions in humans; therefore, carotenoid biofortification of maize (*Zea mays* L.), one of the most highly produced cereal crops worldwide, would have a global impact on human health.
- *PLASTID TERMINAL OXIDASE* (*PTOX*) genes play an important role in carotenoid metabolism; however, the possible function of *PTOX* in carotenoid biosynthesis in maize has not yet been explored. In this study, we identified the maize *PTOX* gene *ZmPTOX1* by forward genetic screening.
- While most higher plant species possess a single copy of the *PTOX* gene, maize carries two tandemly duplicated copies of *ZmPTOX*. Characterization of *Zmptox1* mutants revealed that disrupting one copy of *ZmPTOX1* was enough to impair carotenoid biosynthesis, indicating that *ZmPTOX1* is essential for carotenoid biosynthesis in maize kernels. Remarkably, overexpression of *ZmPTOX1* significantly improved the content of carotenoids, especially β-carotene (provitamin A), which was increased by ~3-fold, in maize kernels.
- Overall, our study shows that *ZmPTOX1* plays a crucial role in carotenoid biosynthesis in maize kernels and suggests that fine-tuning the expression of this gene could improve the nutritional value of cereal grains.

## Introduction

The lack of certain micronutrients poses a serious threat to human health. This is particularly true in developing countries, where people predominantly subsist on cereal grains, which lack certain nutrients. For example, carotenoids, a group of yellow or red pigments with antioxidant activity, are essential for human health but are present at low levels in cereal grains. Therefore, carotenoid biofortification of major cereal crops used as food and feed worldwide would have a global impact on human health (Wurtzel *et al*., 2012; Owens *et al*., 2014).

In plants, carotenoids function as accessory pigments during light harvesting and form integral and structural components of the photosynthetic apparatus. The carotenoid metabolic pathway has been intensively studied in plants (reviewed by (Collini, 2019)). Carotenoids are synthesized in plastids from the colorless pigment phytoene and then subjected to sequential desaturation reactions. The first two desaturation reactions are catalyzed by phytoene desaturase (PDS), which result in the formation of phytofluene and then ζ-carotene (Bartley *et al*., 1991). The ζ-carotene is then catalyzed by ζ-carotene desaturase (ZDS) to form neurosporene and then lycopene (Albrecht *et al*., 1995). Lycopene is cyclized to form either α- or β-carotene, which can be oxidized to produce xanthophylls in photosynthetic tissues. PDS and ZDS require PLASTID TERMINAL OXIDASE (PTOX) for the successive desaturation of carotenoid (Foudree *et al*., 2012).

The first plant *PTOX* gene was identified in the *immutans* (*im*) mutant of *Arabidopsis thaliana* (Carol *et al*., 1999; Wu *et al*., 1999), which show a variegated phenotype. Cells in the green sectors of the *im* mutant possess normal-appearing chloroplasts, while cells in the white sectors lack carotenoid pigments and appear to be blocked at various stages of chloroplast biogenesis (Wetzel *et al*., 1994; Yu *et al*., 2007). In the tomato (*Solanum lycopersicum* L.) *ghost* mutant, orthologous to Arabidopsis *im* mutant, immature fruits appear white because of an overaccumulation of the colorless pigment phytoene and the deficiency of various colored pigments such as lycopene, lutein, and β-carotene (Barr *et al*., 2004). Besides its function in carotenoid biosynthesis, PTOX also regulates plant architecture by affecting the biosynthesis of plant hormones. The rice (*Oryza sativa* L.) *ptox* mutants show excessive tillering, semidwarfism, and variegation because of defects in the biosynthesis of strigolactones (Tamiru *et al*., 2014).

Although the importance of *PTOX* genes for carotenoid metabolism, chloroplast function, and shoot architecture has been well characterized in various plant species, the potential function of these genes in carotenoid biosynthesis in maize (*Zea mays* L.), the most produced cereal crop, has not yet been explored. Here, we identified the maize *PTOX* gene *ZmPTOX1* by forward genetic screening. Characterization of *Zmptox1* mutants established that *ZmPTOX1* plays a critical role in carotenoid biosynthesis in maize kernels. We demonstrate that engineering this gene could enhance the carotenoid content and nutritional value of maize grains.

## Materials and Methods

### Identification and map-based cloning of the *ZmPTOX1* gene

To identify the *PTOX* gene in maize, pollen (2 ml) of the RP125 inbred line (wild type) were treated with 0.066% EMS (Sigma-Aldrich, M0880) solution in mineral oil (Sigma-Aldrich, M8410) for 30 min. The mutagenized pollen was then used to pollinate the ears of RP125. The F2 segregating population was constructed by crossing the M2 *Zmptox1* mutants in the RP125 background with B73. RNA was extracted from ~30 pooled normal or mutant seeds, and BSA-seq was performed as described previously (Gallavotti & Whipple, 2015). Candidate genes in the mapping region were amplified from the mutants and subjected to Sanger sequencing. The deduced amino acid sequences of these candidate genes were aligned using the MUSCLE model, and a phylogenetic tree was constructed using the neighbor-joining method in MEGA7 with 1,000 bootstrap replicates (Kumar *et al*., 2016).

The *ZmPTOX1* overexpression vector was constructed by cloning the coding region of *ZmPTOX1* under the control of the maize ubiquitin promoter. The construct was transformed into *Agrobacterium*, which was then used to transform the maize inbred line B104 as described previously (Frame *et al*., 2015).

### Carotenoid analysis

The carotenoid extraction method described for alfalfa (*Medicago truncatula*) (Meng *et al*., 2019) was used in this study, with minor modifications. Seeds of the following genotypes were used for carotenoid extraction: wild type, *Zmptox1-1, Zmptox1-2, Zmptox1-3*, B104, and OE lines. Briefly, the dissected maize tissues (30 mg) were ground into powder in liquid nitrogen. Then, 200 μL of 6% (w/v) KOH (in methanol) was added to the ground tissue. The samples were vortexed for 10 s and heated at 60°C for 1 h in the dark. Subsequently, 200 μL of 50 mM HCl buffer (pH 7.5, containing 1 M NaCl) was added to each sample. Samples were allowed to cool to room temperature in the dark, thoroughly mixed by turning the tubes upside-down eight to ten times, and then incubated on ice for 10 min. Then, 800 μL of chloroform was added to the sample, mixed by inverting eight to ten times, and incubated on ice for 10 min. The mixtures were centrifuged at 3,000 × g for 5 min at 4°C. Then, the lower liquid phase (600 μL) was removed and again extracted using 800 μL of chloroform. The two chloroform extracts were combined, dried with a blowing device using high quality nitrogen, and then dissolved in 100 mL of ethyl acetate. The different pigments were identified by high performance liquid chromatography (HPLC; A30).

HPLC analysis was performed on endosperm, seed, and other plant tissues as described previously (Fraser *et al*., 2000), with minor modifications. Briefly, a C30 column at 30 ± 1°C was used with solvent A (methanol, methyl cyanide, water, and butylated hydroxytoluene) and solvent B (methyl tert-butyl ether) as mobile phases, with a flow rate of 1.0 mL/min flow rate and an injection volume of 10 μL. Carotenoids were identified by comparing their absorption spectra and retention period with those of the standards. The contents of carotenoids were calculated using standards.

The activities of PDS, ZDS, and PSY enzymes were determined using the ELISA-based method (LMAI Bio), according to the manufacturer’s instructions. Solid-phase plant purified anti-PDS, -ZDS, and -PSY antibodies were used to coat microtiter plate wells. Then, PDS, ZDS, and PSY enzymes were added to the wells. An antibody–antigen–enzyme–antibody complex was labeled with combined antibodies and horse radish peroxidase (HRP). After thorough washing, the reaction was stopped by adding sulfuric acid, which turns the TMB substrate blue when catalyzed by HRP. The color change was then measured spectrophotometrically at a wavelength of 450 nm. The optical density (OD) of the samples was then compared with the standard curve to determine the PDS, ZDS, and PSY concentrations in the samples.

### RNA-seq and data analysis

Kernels of wild-type and *Zmptox1* mutant plants were harvested at 9, 12, and 20 days postpollination (dpp), with three biological replicates at each time point. Total RNA was extracted from the kernels using the TRIzol reagent, as directed by the manufacturer. The sequencing libraries were produced with the Illumina TruSeq RNA Sample Prep Kit and sequenced on the Illumina HiSeq X Ten System. The quality of raw sequence reads was checked using Fastp v.0.12.4 (Chen *et al*., 2018). HISAT2 v.2.2.1, with default settings, was used to map the clean reads to the maize B73 RefGen_V4 reference genome sequence (Kim *et al*., 2015). StringTie v.2.0.4 was utilized to calculate gene expression levels as fragments per kilobase of transcript per million reads (FPKM) (Pertea *et al*., 2016). DEGs were identified using DESeq2 based on two criteria: log2fold-change ≥1 and adjusted *p*-value cutoff ≤0.05 (Love *et al*., 2014).

### Candidate gene association and nucleotide diversity analyses

Candidate gene association analysis was conducted using a maize association panel of 368 diverse inbred lines, and *ZmPTOX1*-specific SNPs and metabolite content data of the members of this panel were downloaded from the previously released genotype dataset (Fu *et al*., 2013). Association between metabolite contents and *ZmPTOX1*-specific SNPs was determined using a mixed linear model, after correction for population structure, with a *p*-value of 0.001 as a threshold (Li *et al*., 2013). The third generation haplotype map data of *Zea mays* were downloaded. Nucleotide diversity was investigated in improved maize lines, landraces, and teosinte using vcftools (https://vcftools.github.io/examples.html), with a 1 kb window and 100 bp step along the 2 kb upstream and 1 kb downstream regions of *ZmPTOX1*, respectively.

## Results

### Map-based cloning of *ZmPTOX1*

To identify the key genes involved in carotenoid biosynthesis in maize kernels, we performed ethyl methanesulfonate (EMS) mutagenesis screening on the Chinese elite maize inbred line RP125 and searched for mutants with pale kernels, an indicator of low carotenoid accumulation in the endosperm (Nie *et al*., 2021). A mutant with much paler kernels than the wild type (RP125, yellowish kernels) was identified in the M2 population (Figure 1a). The mutants also displayed leaf variegation, with white and green sectors, at the seedling stage (Figure 1b) and accumulated less chlorophyll than the wild-type plants (Figure 1c). Together, these results suggest that the mutants exhibit not only low carotenoid accumulation but also abnormal chloroplast biogenesis.

**Figure 1.**
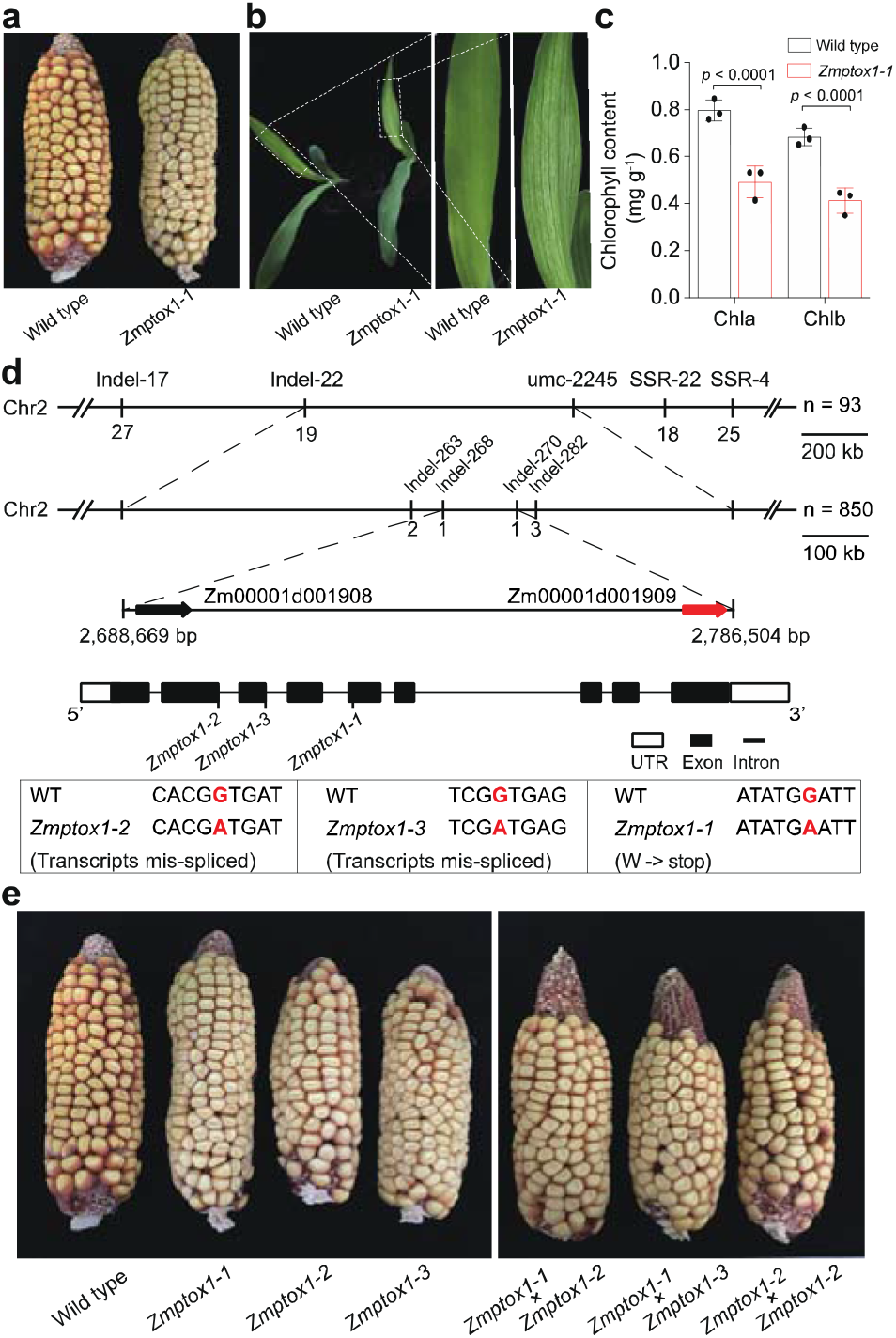
Map-based cloning of *ZmPTOX1* (a) Ear phenotypes of the wild-type (RP125) and *Zmptox1* mutants. The kernels of mutants are paler than those of the wild type. (b) Leaf phenotype of wild-type and *Zmptox1* mutant plants. The leaves of mutants are paler than those of the wild type. (c) Chlorophyll content of the *Zmptox1* mutant was significantly higher than that of the wild type. (d) Scheme for positional cloning of the *Zmptox1* gene. (e) Comparison of the ear phenotypes of allelic mutants and their progenies with that of the wild type.

To identify and clone the causal gene, we generated a recombinant F2 population by crossing the recessive mutant with B73 inbred line and performed bulked segregant analysis (BSA) using pooled RNA samples representing approximately 30 mutant or wild-type kernels (Gallavotti & Whipple, 2015). We mapped the gene to an approximately 2 Mb region on the short arm of chromosome 2. Using ~850 individual mutants, we narrowed down the region to ~100 kb containing two putative genes, *Zm00001d001908* and *Zm00001d001909*. Sequencing revealed a G-to-A mutation in the exon 5 of *Zm00001d001909*, which was predicted to change the tryptophan residue at position 186 to a premature stop codon (Figure 1d). We also identified two additional mutants, with similar phenotypes, in our EMS collection. Both mutants carried allelic mutations at splice sites in *Zm00001d001909*, leading to incorrectly spliced transcripts (Figure 1d). The F1 plants obtained by crossing these mutants displayed similar phenotypes (Figure 1e), confirming that the two mutants were allelic, and that the mutations in *Zm00001d001909* were responsible for the pale phenotype of kernels.

### *ZmPTOX1* regulates carotenoid biosynthesis in maize kernels

Protein BLAST and phylogenetic analysis revealed that *Zm00001d001909* encodes PTOX (Figure 2a), an enzyme that activates the plastid enzyme PDS, which in turn converts the colorless phytoene to ζ-carotene, a precursor of colored carotenoids ((Wetzel *et al*., 1994) and Figure 2b). PTOX acts as a cofactor of PDS and ZDS, two enzymes required for carotenoid biosynthesis. To determine whether this mechanism is conserved in maize kernels, we examined the activities of PDS and ZDS in wild-type and *Zmptox1-1* kernels. Both enzymes showed reduced activity in mutant kernels compared with wild-type kernels (Figure 2c and d), suggesting that ZmPTOX1 participates in carotenoid biosynthesis by regulating the activity of PDS and ZDS.

**Figure 2.**
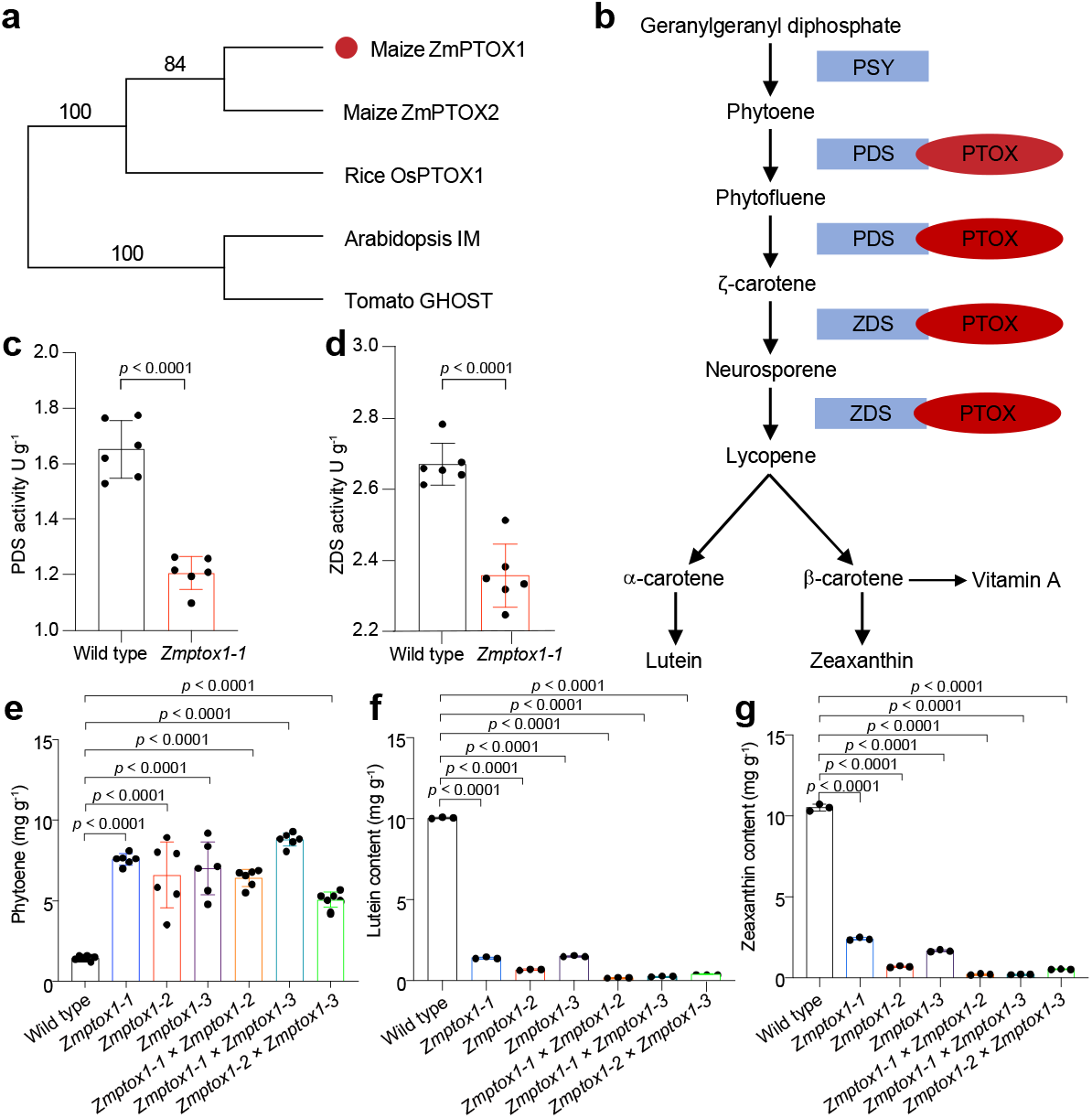
*ZmPTOX1* regulates carotenoid biosynthesis in maize kernels. (a) Phylogenetic analysis of *PTOX* genes of maize, rice, Arabidopsis, and tomato. (b) Biosynthetic pathway of lutein and zeaxanthin. (c, d) Activities of PDS (c) and ZDS (d) in *Zmptox1* mutants and wild-type (RP125) plants. (e–g) Contents of phytoene (e), lutein (f), and zeaxanthin (g) in wild-type and *Zmptox1* kernels.

To confirm whether the pale color of mutant kernels was indeed caused by the lower accumulation of colored carotenoids, we measured the contents of colored carotenoids, including lutein, zeaxanthin, and carotene, as well as that of phytoene in wild-type and mutant kernels. The mutant kernels accumulated more noncolored pigments such as phytoene (Figure 2e) and fewer colored pigments such as lutein and zeaxanthin (Figure 2f and g). Additionally, neither α-carotene nor β-carotene was detected in mutant kernels. These results confirmed that the pale color of mutant kernels was caused by the disruption of *ZmPTOX1*, indicating that the loss of *ZmPTOX1* impairs carotenoid biosynthesis in maize.

To gain further insights into the transcriptomic changes in *Zmptox1* mutant kernels, we performed RNA-seq analyses on wild-type and *Zmptox1-1* mutant kernels sampled at 9, 12, and 20 dpp. Principal component analysis (PCA) showed strong correlation among the biological replicates of each genotype, indicating that our RNA-seq data were highly reliable (Supplementary Figure 1a). Comparison between the RNA-seq data of *Zmptox1* and wild-type kernels led to the detection of 1,523 (905 upregulated and 618 downregulated), 571 (283 upregulated and 288 downregulated), and 523 (277 upregulated and 246 downregulated) differentially expressed genes (DEGs) at 9, 12, and 20 dpp, respectively (Supplementary Table 1). A total of 26 DEGs were significantly upregulated, and 26 DEGs were downregulated, at least at one time point (Supplementary Figure 1b and c). Functional enrichment analysis of the DEGs using the Kyoto Encyclopedia of Genes and Genomes (KEGG) database revealed 20 representative pathways at three developmental stages (Figure 3a), among which the ‘plant-pathogen interaction pathway’ was the most highly enriched, suggesting that either PTOX functions in pathogen resistance or changes in carotenoid metabolism affect pathogen fitness. A closer look at the transcriptomes revealed that the expression of genes involved in methylerythritol phosphate (MEP), mevalonate (MVA), and carotenoid pathways remained largely unchanged in *ptox1* mutant kernels compared with wild-type kernels (Figure 3b). These results were anticipated, given that *PTOX1* is not a regulatory gene.

**Figure 3.**
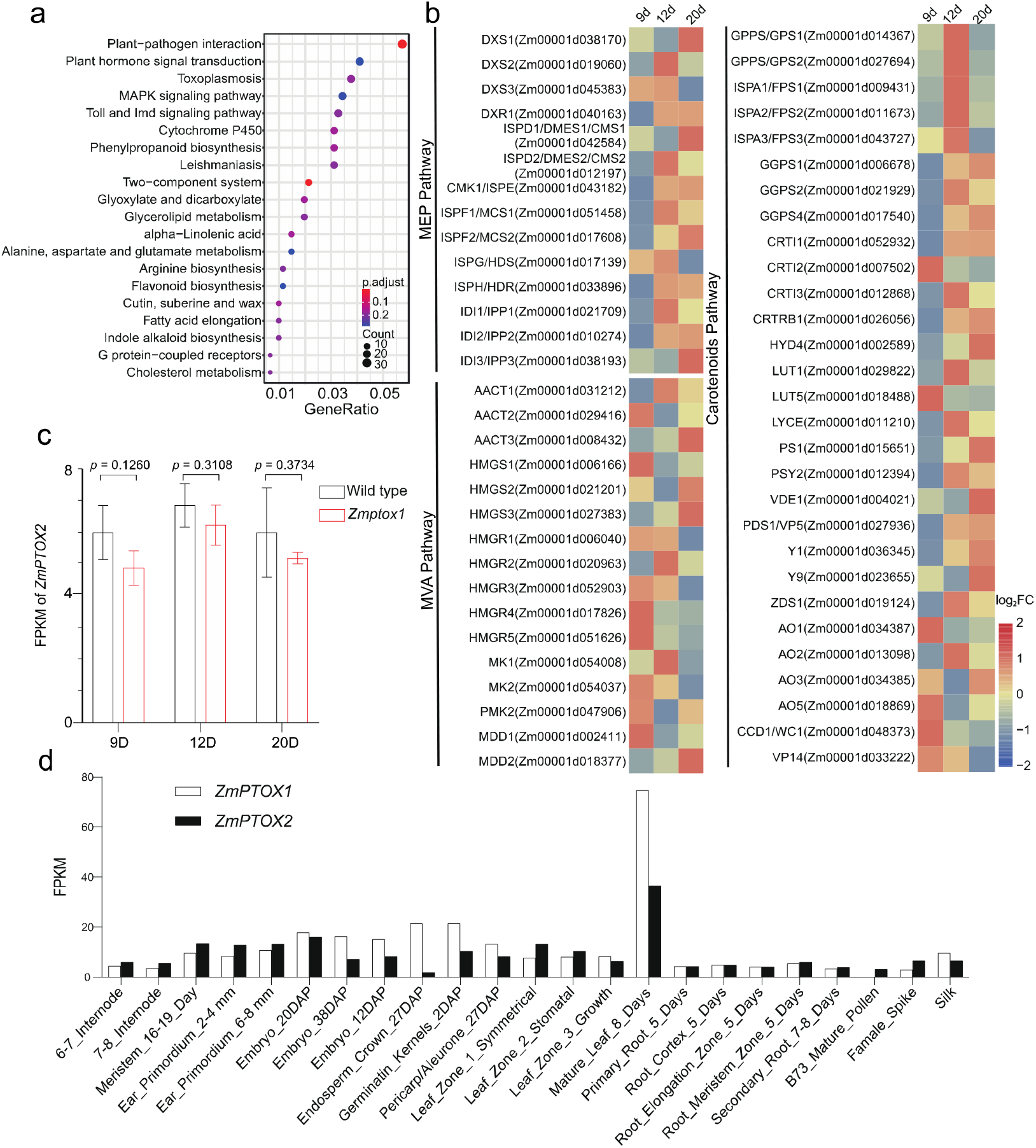
Transcript profiling of wild-type and *Zmptox1* mutant kernels at 9, 12, and 20 dpp. (a) KEGG enrichment of DEGs. (b) Comparison of the expression levels of MEP, MVP, and carotenoid pathway genes in wild-type controls and *Zmptox1* mutants. (c, d) Expression levels of *ZmPTOX1* and *ZmPTOX2* in mutant and wild-type kernels (c) and other tissues (d).

Phenotypes associated with the disruption of *PTOX* have been reported in other plant species such as Arabidopsis, tomato, and rice; however, these phenotypes were related only to changes in leaf color, fruit color, and shoot architecture and not to changes in seed color (Carol *et al*., 1999; Wu *et al*., 1999; Barr *et al*., 2004; Tamiru *et al*., 2014). Additionally, Arabidopsis, tomato, and rice possess only one copy of *PTOX*, whereas maize has two tandemly duplicated *PTOX* genes separated by ~80 kb (Figure 2a). Accordingly, we designated *Zm00001d001909* as *ZmPTOX1* and its paralog *Zm00001d001908* as *ZmPTOX2. ZmPTOX1* transcripts were more abundant than *ZmPTOX2* transcripts (Figure 3d), consistent with a previous study (Walley *et al*., 2016)), which explains why the impairment of *ZmPTOX1* alone was sufficient to cause pale kernels and variegated leaves. However, unlike some other paralogous genes whose expression is upregulated to complement the nonfunctional gene paralog in the single mutant (Rodriguez-Leal *et al*., 2019), the expression of *ZmPTOX2* was unchanged in *Zmptox1* mutants, explaining why the ablation of a single *ZmPTOX* gene was sufficient to affect carotenoid metabolism (Figure 3c).

To determine whether natural variation in *ZmPTOX1* is associated with carotenoid content, we conducted a candidate gene association study using a maize association panel (*n* = 368 diverse inbred lines) and previously published carotenoid content data (Fu *et al*., 2013). The results revealed five single nucleotide polymorphisms (SNPs) in *ZmPTOX1*, all of which showed significant association with carotenoid content. Three of these SNPs were associated with α-carotene content, while the remaining two SNPs were associated with the lutein content of kernels (Figure 4a). The two SNPs associated with lutein content were found in the promoter region of *ZmPTOX1*. Among the three α-carotene content-associated SNPs, two were found in the intron and one in the exon (2,779,620 bp). The exonic SNP was nonsynonymous and predicted to cause proline-to-serine amino acid substitution. To further explore the effect of the exonic SNP on carotenoid content, it would be interesting to examine if the amino acid substitution directly alters the enzyme activity of ZmPTOX1. Overall, our findings showed that natural variation in *ZmPTOX1* influences lutein and α-carotene contents (4b, c), suggesting that the manipulation of this gene may enhance the carotenoid content of maize kernels.

**Figure 4.**
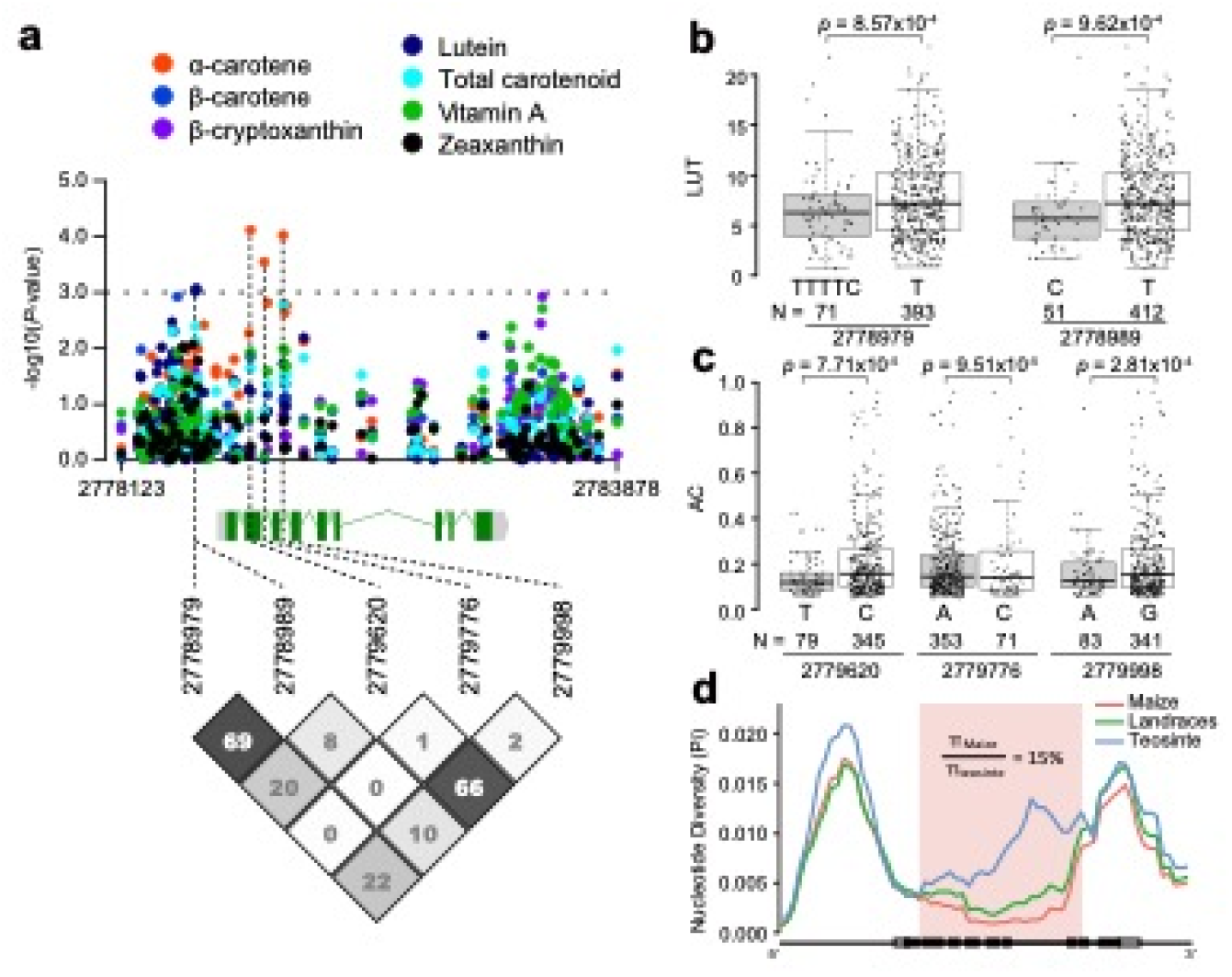
Natural variation in the promoter and open reading frame (ORF) of *ZmPTOX1* is associated with the pale color of maize kernels. (a) Two SNPs in the promoter and three SNPs in the ORF of *ZmPTOX1* show association with lutein (LUT) and α-carotene (AC) contents, respectively. The dashed line indicates the threshold of significant association (*p* ≤ 1.00E-03). The numbers show the pairwise linkage disequilibrium pattern by R^2^ (%). The positions of five significant SNPs are shown on the top of the pairwise linkage disequilibrium plot and indicated by dashed lines in the gene diagram. (b, c) The contents of lutein (LUT) and a-carotene (AC) associated with SNPs in diverse inbred lines. Each box represents the median and interquartile range, and whiskers extend to maximum and minimum values. The genotype and N of each allele are listed below each box. (d) Evidence of selection pressure on *ZmPTOX1*. The red area shows low nucleotide diversity in improved maize lines compared with teosinte, as analyzed using maize HapMap v3 SNP data. Red, green, and blue lines represent the nucleotide diversity of improved maize lines, landraces, and teosinte, respectively. White and black rectangles on the x-axis represent the untranslated regions (UTRs) and exons of the *ZmPTOX1* gene.

A previous study showed that maize represents, on average, only 57.1% of the nucleotide diversity in teosinte (Wright *et al*., 2005). To determine whether *ZmPTOX1* underwent selection during maize domestication and improvement, we calculated the nucleotide diversity of *ZmPTOX1* in improved maize lines, landraces, and teosinte using HapMap 3 (Bukowski *et al*., 2017). Nucleotide diversity was high in teosinte but significantly reduced in the improved maize lines, especially in the region highlighted in red in Figure 4d. These data suggest that *ZmPTOX1* underwent positive selection throughout the maize improvement process; however, the implication of this selection remains to be explored.

### Overexpression of *ZmPTOX1* improves nutritional content of maize kernels

To determine whether *ZmPTOX1* could directly enhance the nutritional content of maize kernels, we overexpressed *ZmPTOX1* under the control of the maize ubiquitin promoter. Ten independent events were obtained, three of which were randomly selected for further analysis. Quantitative real-time PCR (qRT-PCR) revealed that *ZmPTOX1* was upregulated 10–45-fold in all three overexpression (OE) lines relative to the wild-type controls (Figure 5a). Additionally, the levels of lutein, zeaxanthin, and α-carotene were nearly 2-fold greater, and the levels of the β-carotene were 3-fold higher, in transgenic kernels than in wild-type controls (Figure 5b–e). These findings demonstrate that manipulating *ZmPTOX1* expression is effective in improving maize carotenoid content.

**Figure 5.**
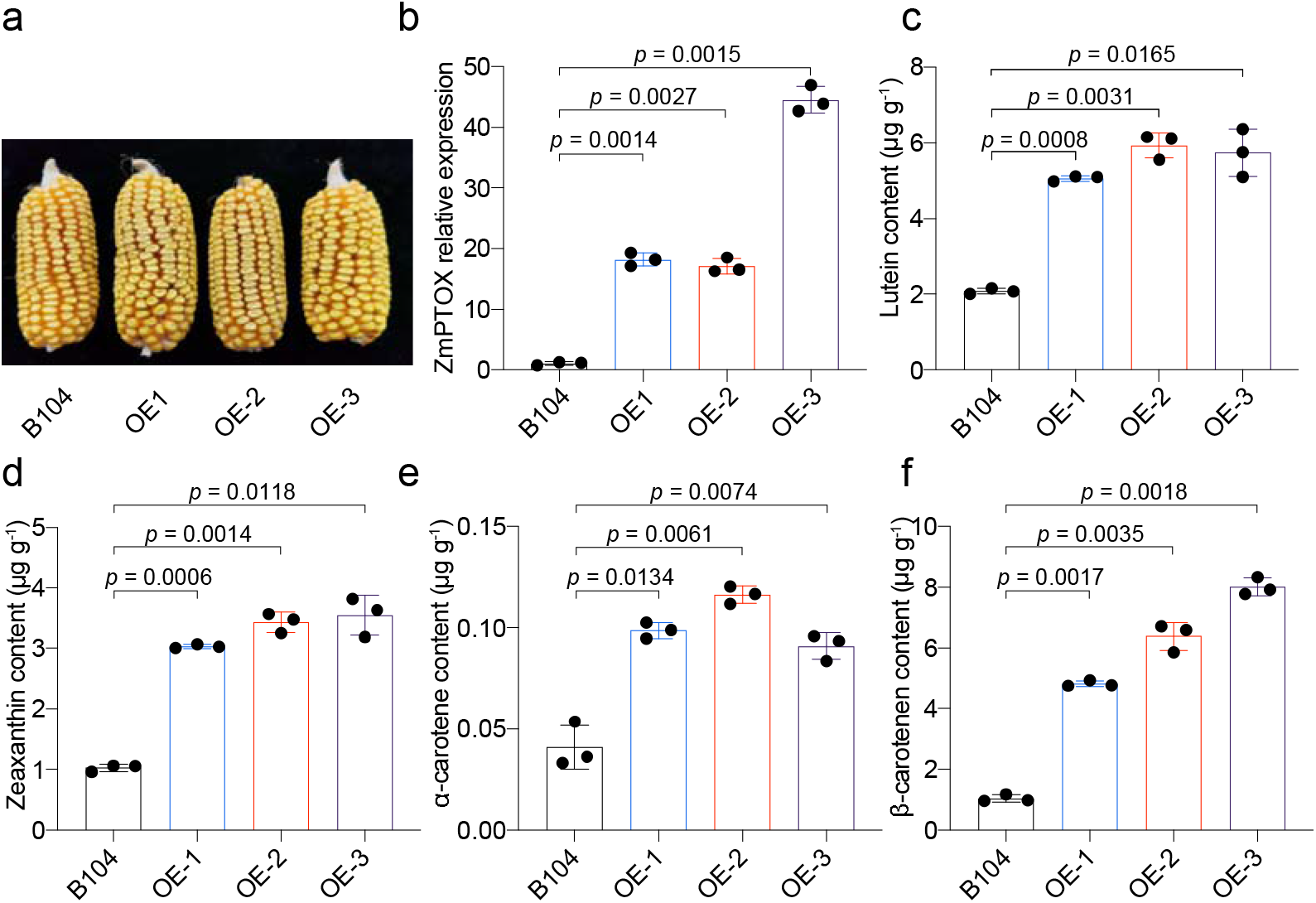
Characteristics of three *ZmPTOX1*-overexpressing maize lines (OE1–3). (a) Ears of wild-type (B104), OE1, OE2, and OE3 plants. (b) *ZmPTOX1* transcripts were more abundant in the transgenic lines. (c–f) Overexpression of *ZmPTOX1* increased the contents of lutein (c), zeaxanthin (d), α-carotene (e), and β-carotene (f). In (b–f), data represent mean ± standard deviation (SD; *n* = 3). Significant differences were determined using Student’s *t*-test. Raw data are plotted in the bar charts.

## DISCUSSION

Through forward genetic screening, we identified a *PTOX* gene in maize, one of the most important cereal crops worldwide, and demonstrated that engineering this gene dramatically increases the levels of carotenoids, particularly β-carotene, a nutrient crucial for human health. The carotenoid biosynthetic pathway in plants has been established through the cloning and characterization of genes using specific carotenoid mutants. Although phytoene synthase (PSY), which is responsible for phytoene biosynthesis, has received much attention as a rate-limiting enzyme in this pathway (Zhou *et al*., 2015; Chayut *et al*., 2017), the PDS and ZDS enzymes, which desaturate phytoene and ζ-carotene, are also critical for the biosynthesis of downstream health-promoting carotenoids (McQuinn et al). In tomato, PDS becomes limiting once PSY1 is elevated during fruit ripening (McQuinn *et al*., 2018). Thus, new bottlenecks emerge in subsequent desaturation steps to promote the activity of PDS and ZDS. Our data show that PTOX, the cofactor of PDS and ZDS, is a great target for engineering maize cultivars with high carotenoid content.

Although the maize genome carries two tandemly duplicated *PTOX* genes, in contrast to the single *PTOX* gene in *Arabidopsis*, tomato, and rice, disrupting one of the two copies in maize is sufficient to impair carotenoid biosynthesis in grains. *PTOX* exists as a single-copy gene in most plant species (Carol *et al*., 1999; Wu *et al*., 1999; Barr *et al*., 2004; Tamiru *et al*., 2014); however, in some cyanobacteria and algae, two copies of the *PTOX* gene have been reported (Wang *et al*., 2009; Houille-Vernes *et al*., 2011), of which one copy is more important than the other. For example, *Chlamydomonas* has *PTOX1* and *PTOX2;* however, PTOX2 is the predominant enzyme isoform involved in chlororespiration, and the *ptox2* single mutant shows lower fitness than the wild type when grown under phototrophic conditions (Houille-Vernes *et al*., 2011). Our study is the first to show that higher plants contain two copies of *PTOX*, and that disrupting one copy is sufficient to cause obvious phenotypes. Although both *ZmPTOX1* and *ZmPTOX2* were expressed in a variety of tissues, *ZmPTOX1* transcripts were more abundant than *ZmPTOX2* transcripts in the majority of maize tissues (Figure 3d and (Walley *et al*., 2016)), particularly mature leaves and endosperm, which showed clear changes in color because of decreased pigment accumulation (Figure 1a,c; Figure 2h). Our findings imply that *ZmPTOX1* plays a major role in carotenoid biosynthesis, and its disruption is sufficient to impair this process. However, whether the maize *PTOX* genes exhibit a dosage effect remains to be explored. Elevating the abundance of *ZmPTOX1* transcripts improved downstream carotenoids, indicating that dosage effect is likely (Figure 5). However, detailed characterization of *Zmptox1* and *Zmptox2* single and double mutants is required before drawing any firm conclusions. Unfortunately, the two genes are only ~80 kb apart. Therefore, creating double mutants by crossing the single mutants and screening for recombinants would be highly challenging. To overcome this issue, the CRISPR-Cas system could be used to knockout both genes simultaneously.

In addition to chloroplast biogenesis and carotenoid biosynthesis, PTOXs are potentially also involved in other physiological processes such as stress tolerance and plant architecture regulation (Tamiru *et al*., 2014; Johnson & Stepien, 2016). For example, transgenic tobacco plants transformed with the *PTOX1* gene of the green alga *Chlamydomonas reinhardtii* outperformed wild-type controls in terms of seed germination rate, root length, and shoot biomass accumulation under high NaCl concentration. Transgenic tobacco plants also displayed better recovery and less chlorophyll bleaching than wild-type plants after the NaCl treatment (Ahmad *et al*., 2020). Rice *ptox* mutants displayed excessive tillering and a semidwarf phenotype, demonstrating that PTOX functions in strigolactone biosynthesis (Tamiru *et al*., 2014). However, the *Zmptox1* mutants did not exhibit any developmental phenotypes compared with wild-type plants. One explanation is that the two *ZmPTOX* genes in maize perform redundant functions; thus, *ZmPTOX2* can compensate for the loss of function of *ZmPTOX1* and is adequate to sustain the biosynthesis of strigolactones in the *Zmptox1* mutant. To verify whether *ZmPTOX1* and *ZmPTOX2* are redundant genes, a thorough analysis of the developmental characteristics of *Zmptox* double mutants is needed. Given the diverse functions of PTOXs in regulating carotenoid biosynthesis, plant development, and stress resistance, fine-tuning the expression of *ZmPTOX1* has a great potential to improve the nutritional value, optimize the plant architecture, and enhance the stress tolerance of maize simultaneously.

## Acknowledgements

Funding for this work was provided by the National Natural Science Foundation of China (32171925, U21A20210) to Q.W.; The National Key Research and Development Program of China (2022YFD1201703), Taishan Scholars Program of Shandong Province (tsqn201909074), and Shandong Natural Science Foundation (ZR2020KC019) to Z.Z.; The Agricultural Science and Technology Innovation Program(ASTIP) to Q.W.; Key R&D Program of Shandong Province (2021LZGC022-2) to H.D..; Shandong Natural Science Foundation (ZR2022MC076) to Y.N.; and National Science Foundation (NSF) to DJ (IOS-2129189).

## Author contributions

Q.W., Z.Z., Y.N., H.W., G.Z., H.D, D.J, G.P, X.Y.Z., X.S.Z., and X.H.L. designed research and wrote the paper. Y.N., H.W., G.Z., and H.D. cloned the *ZmPTOX1* gene. J.S., J.D., X.L., X.Z.L., Y.Z., X.Z, C.L., J.W., G.PZ.Q., K.Z., Q.L., Y.C., C.Z., C.L., J.W., X.L., X.S.Z., X.Y.Z, G.P., and B.H. generated the transgenic plants and determined the carotenoid contents. L.L. and D.J. performed candidate gene association analysis.

## Data availability

The raw data are available from Zhiming Zhang or Qingyu Wu upon request. All RNAseq datasets reported in this study have been deposited in GenBank (NCBI) with the following accession code: PRJNA943556.

---

**Supplementary Figure 1.**
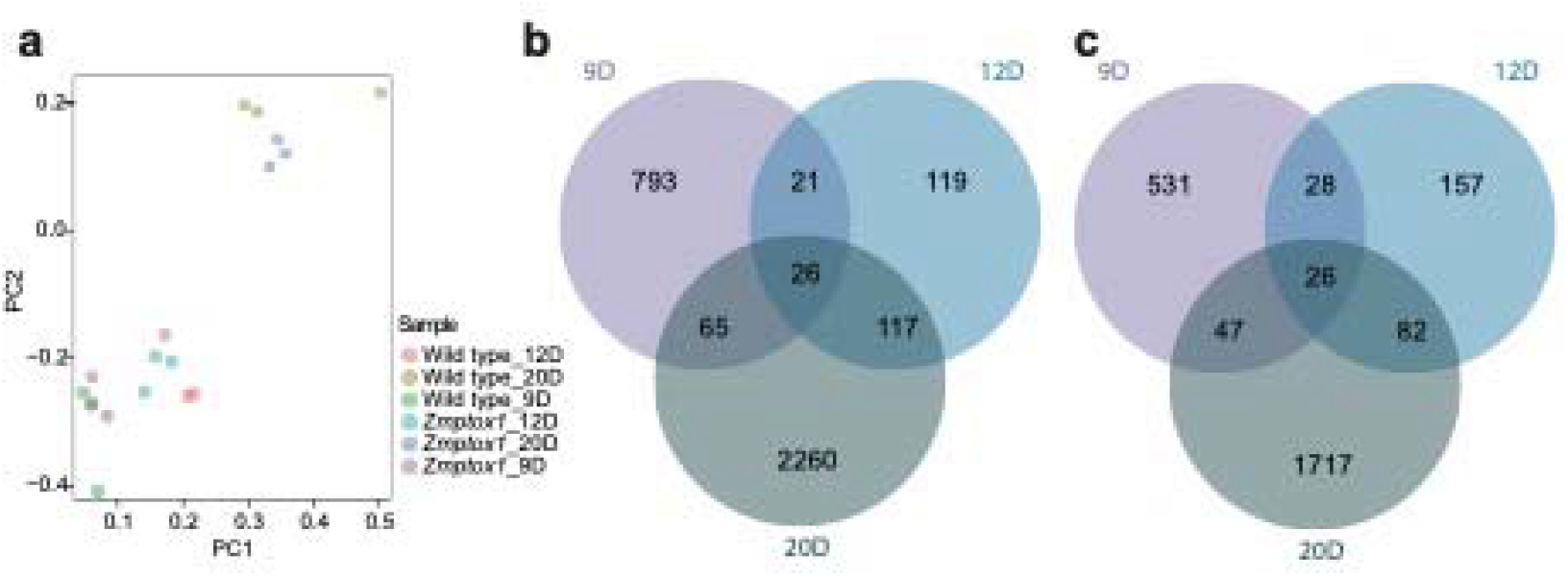
Statistics of DEGs. (a) Principal component analysis (PCA) of transcriptome data. (b and c) Venn diagram of DEGs at 9, 12 and 20d after pollination.

**Supplementary Table 1** The list of differentially expressed genes.

