## Supplementary figures and images for "The maize *PLASTID TERMINAL OXIDASE* (*PTOX*) gene controls the carotenoid content of kernels"

### Supplementary Figure 1

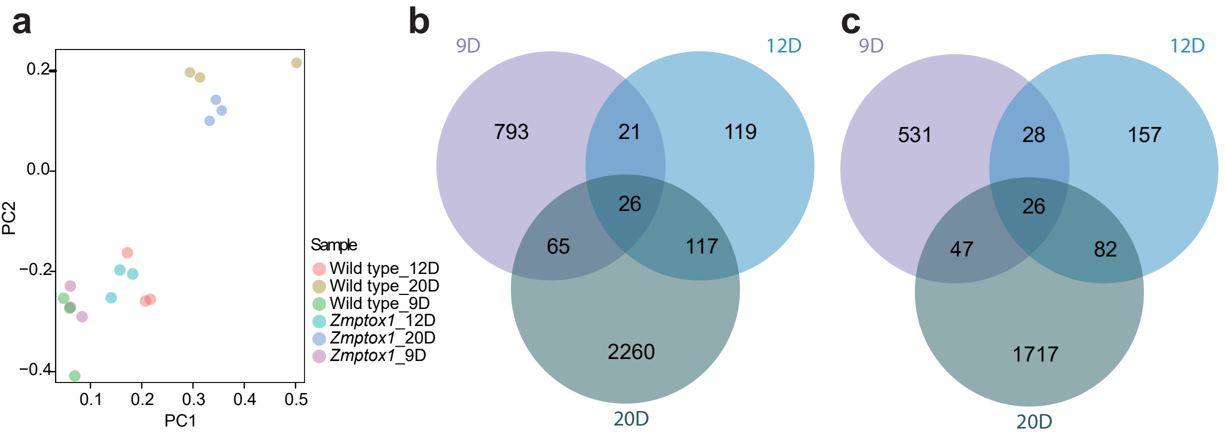
